# Deep-Channel uses deep neural networks to detect single-molecule events from patch-clamp data

**DOI:** 10.1101/767418

**Authors:** Numan Celik, Fiona O’Brien, Sean Brennan, Richard D. Rainbow, Caroline Dart, Yalin Zheng, Frans Coenen, Richard Barrett-Jolley

## Abstract

Single-molecule research such as patch-clamp recording delivers unique biological insight by capturing the movement of individual proteins in real time, unobscured by whole-cell ensemble averaging. The critical first step in analysis is event detection, so called “idealisation”, where noisy raw data are turned into discrete records of protein movement. To date there have been practical limitations in patch-clamp data idealisation; *high quality* idealisation is typically laborious and becomes infeasible and subjective with complex biological data containing many distinct native single ion channel proteins gating simultaneously. Here we show a deep learning model based on convolutional neural networks and long short-term memory architecture can automatically idealise complex single molecule activity more accurately and faster than traditional methods. There are no parameters to set; baseline, channel amplitude or numbers of channels for example. We believe this approach could revolutionise the unsupervised automatic detection of single-molecule transition events in the future.

## Introduction

Ion channels produce functional data in the form of electrical currents typically recorded with the Nobel Prize winning patch-clamp electrophysiological technique^1, 2^. The role of ion channels in the generation of the nerve action potentials was first described in detail in the Nobel Prize winning work of Hodgkin and Huxley^3^, but it is now known they sub-serve a wide range of processes via control of the membrane potential^4^. Loss or dysregulation of ion channels directly underlies many human and non-human animal diseases (so called channelopathies); including cardiovascular diseases such as LQT associated Sudden Death^5^. The first step in analysing ion channel or other single molecule data (which may, in fact, include several individual “single” proteins) is to idealise the noisy raw data. This is typically accomplished by human supervised threshold-crossing although other human supervised methods are available^6, 7^. This produces time-series data with each time point binary classified as open or closed; with more complex data this is a categorical classification problem, with classifiers from zero to *n* channels open. Similar data are also acquired from other single molecule techniques such as lipid bilayer^8^ or single molecule FRET^9–11^. These data can then be used to re-construct the hidden Markov stochastic models underlying the protein activity, using applications such as HJCFIT^12^, QuB (SUNY, Buffalo^13^) or SPARTAN^11^. The initial idealisation step, however, is well recognised by electrophysiologists as a time consuming and labour-intensive bottleneck. This was perhaps best summarised by Professors Sivilotti and Colquhoun FRS^14^ “[patch-clamp recording is] *the oldest of the single molecule techniques, but it **remains unsurpassed [in] the time resolution that can be achieved***. *It is the richness of information in these data that allows us to study the behaviour of ion channels at a level of detail that is unique among proteins. [BUT] This quality of information comes at a price* […]. ***Kinetic analysis is slow and laborious***, *and its success cannot be guaranteed, even for channels with good signals”*. In the current report we show that the solution to these problems could be to apply the latest deep learning methodology to single channel patch-clamp data analyses. For straightforward research with *manual patch-clamp equipment*, and patches with only one or two channels active at a time, it could be argued that the current methods are satisfactory, however, from our own experience, many patches have several channels gating simultaneously and need to be discarded, wasting experimenter time and quite possibly, increasing the numbers of animal donors required. Furthermore, several companies have now developed automated, massively parallel, “patch-clamp” machines^15^ that have the capacity to generate dozens or even hundreds of simultaneous recordings. Use of this technology for single channel recording is greatly compromised by limitations with current software. For example, in most currently available solutions the user has to set the (i) the number of channels in the patch, (ii) the baseline and (iii) the size of the channel manually. If there is baseline drift this would need to be corrected to achieve acceptable results. Our vision is that new deep learning methodology, could in the future make such analyses entirely plausible.

Deep learning^16^ is a machine learning development that has been used to extract features and/or detect objects from different types of datasets for classification problems including base-calling in single-molecule analysis^17, 18^. Convolutional neural network (CNN) layers are a powerful component of deep learning useful for learning patterns within complex data. 2-dimensional (2D) CNNs are most commonly applied to computer vision^19–21^ and we have previously used them for automatic diagnosis of retinal disease in images^22^. An adaptation of the 2D CNN is the one-dimensional (1D) CNN. These have been specifically developed to bring the power of the 2/3D CNN to frame-level classification of time series, and have previously been used in nanopore time-series single-molecule event classification^23^, but never previously patch clamp data. More commonly, the deep learning architecture known as recurrent neural networks (RNNs) have been applied to time series analyses^24, 25^. General RNNs are a useful model for text/speech classification and object detection in time series data, however the model begins degrading once output information depends on long time scales due to a vanishing gradient problem^26^. Long short-term memory (LSTM) networks, are a type of RNN that resolve this problem^27–29^. While 1D-CNN layers can effectively classify raw sequence data, in the current work we combine these with LSTM units to improve the detection of learn long term temporal relationships in time series data^30^.

In the current work, we introduce a hybrid recurrent convolutional neural network (RCNN) model to idealise ion channel records, with up to 5 ion channel events occurring simultaneously. To train and validate models, we developed an analogue synthetic ion channel record generator system and find that our Deep-Channel model, involving LSTM and CNN layers, rapidly and accurately idealises/detects experimentally observed single molecule events without need for human supervision. To our knowledge, this work is the first deep learning model designed for the idealisation of patch-clamp single molecule events.

## Results

### Benchmarking Deep-Channel event detection against human supervised analyses

Our data generation workflow is illustrated in Fig. 1a,c and our Deep-Channel architecture in Fig. 2. In training and model development we found that whilst LSTM models gave good performance, the combination with a time distributed CNN gave increased performance (Supplementary data 1), a so called RCNN we call here Deep-Channel. After training and model development (see methods) we used 17 newly generated datasets, previously unseen by Deep-Channel, and thus uninvolved with the training process. Authentic ion channel data (Fig. 1b) were generated as described in the methods from two kinetic schemes, the first; M1 (see methods and Fig. 3a) with low channel open probability, and the second; M2 with a high open channel probability and thus an average of approximately 3 channels open at a time (Fig. 3b). Across the datasets we included data from both noisy, difficult to analyse signals and low noise (high signal to ratio samples) as would be the case in any patch-clamp project. Examples of these data, together with ground truth and Deep-Channel idealisation are shown in Fig. 4. Note that all the Deep-Channel results described in this manuscript were achieved with a single deep learning model script [capacity to detect a maximum of 5 channels] with no human intervention required beyond giving the script the correct filename/path. So, to clarify; there was no need to set baseline, channel amplitude or number of channels present etc.

**Fig. 1.**
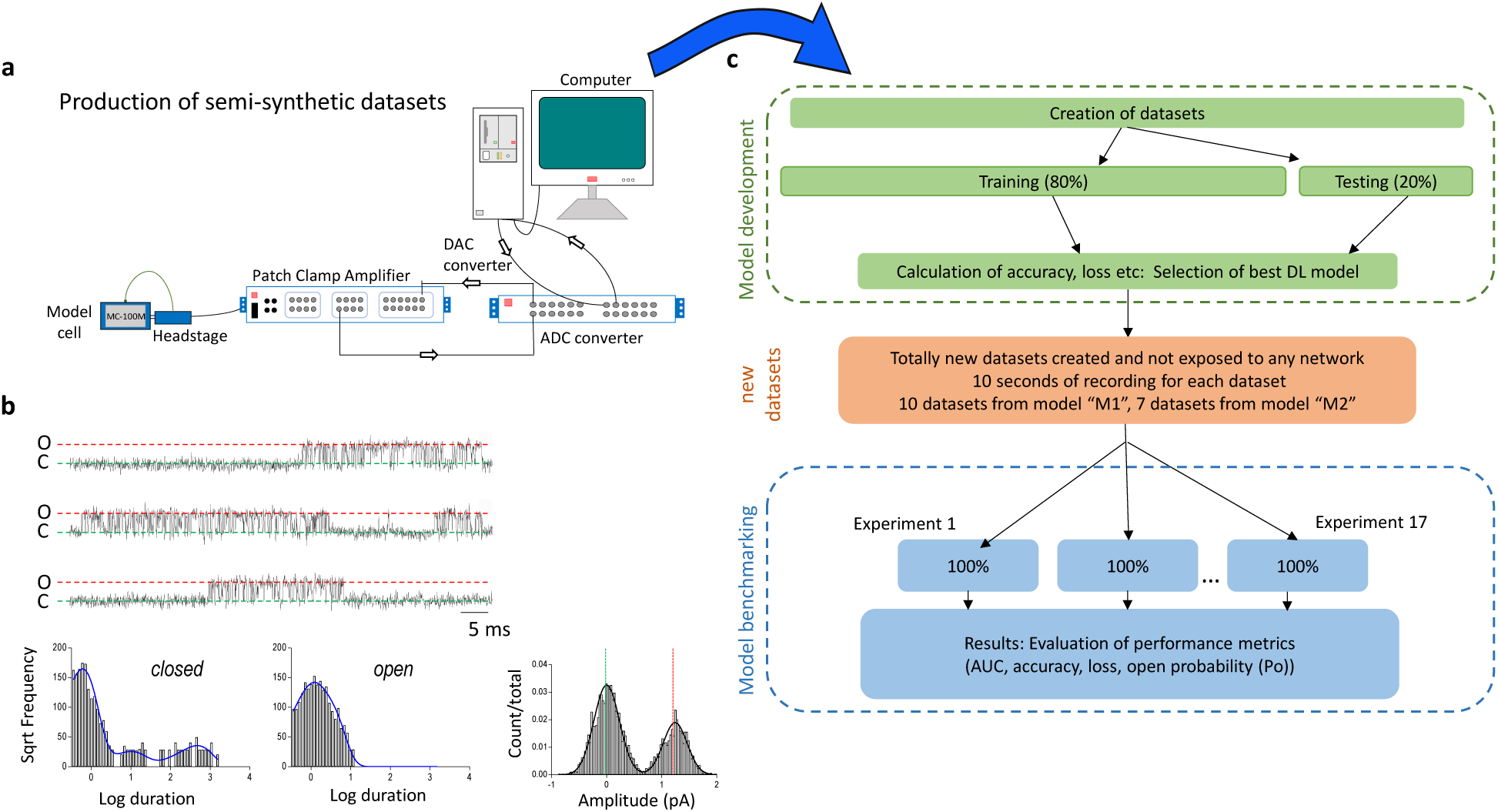
Workflow diagram: Generation of artificial analogue datasets. **a,** For both training, validation and benchmarking, data were generated first as fiducial records with authentic kinetic models in MATLAB (Fig. 2); these data were then played out through a CED digital to analogue converter to a patch clamp amplifier that sent this signal to a model cell and recorded the signal back (simultaneously) to a hard disk with CED Signal software via a CED analogue to digital converter. The degree of noise could be altered simply by moving the patch-clamp headstage closer or further to the PC. In some cases, drift was added as an additional challenge via a separate Matlab script. **b,** Raw single channel patch clamp data produced by these methods are visually indistinguishable from genuine patch clamp data. To illustrate this point, we show here a standard analysis work-up for one such experiment with **bi,** raw data, then it’s analyses with QuB: kinetic analyses of **bii** channel open and **biii** closed dwell times. Finally we show **biv,** all points amplitude histogram. The difference between this and standard ion channel data is that here we have a perfect fiducial record with each experimental dataset, which is impossible to acquire without simulation. **c,** Illustrates our over-all model design and testing workflow. The supplementary data includes training metrics from the initial validation and the main text here shows performance metrics acquired from 17 experiments with entirely new datasets. The training datasets typically contained millions of sample points and the 17 benchmarking experiments were sequences of 100,000 samples each.

**Fig. 2.**
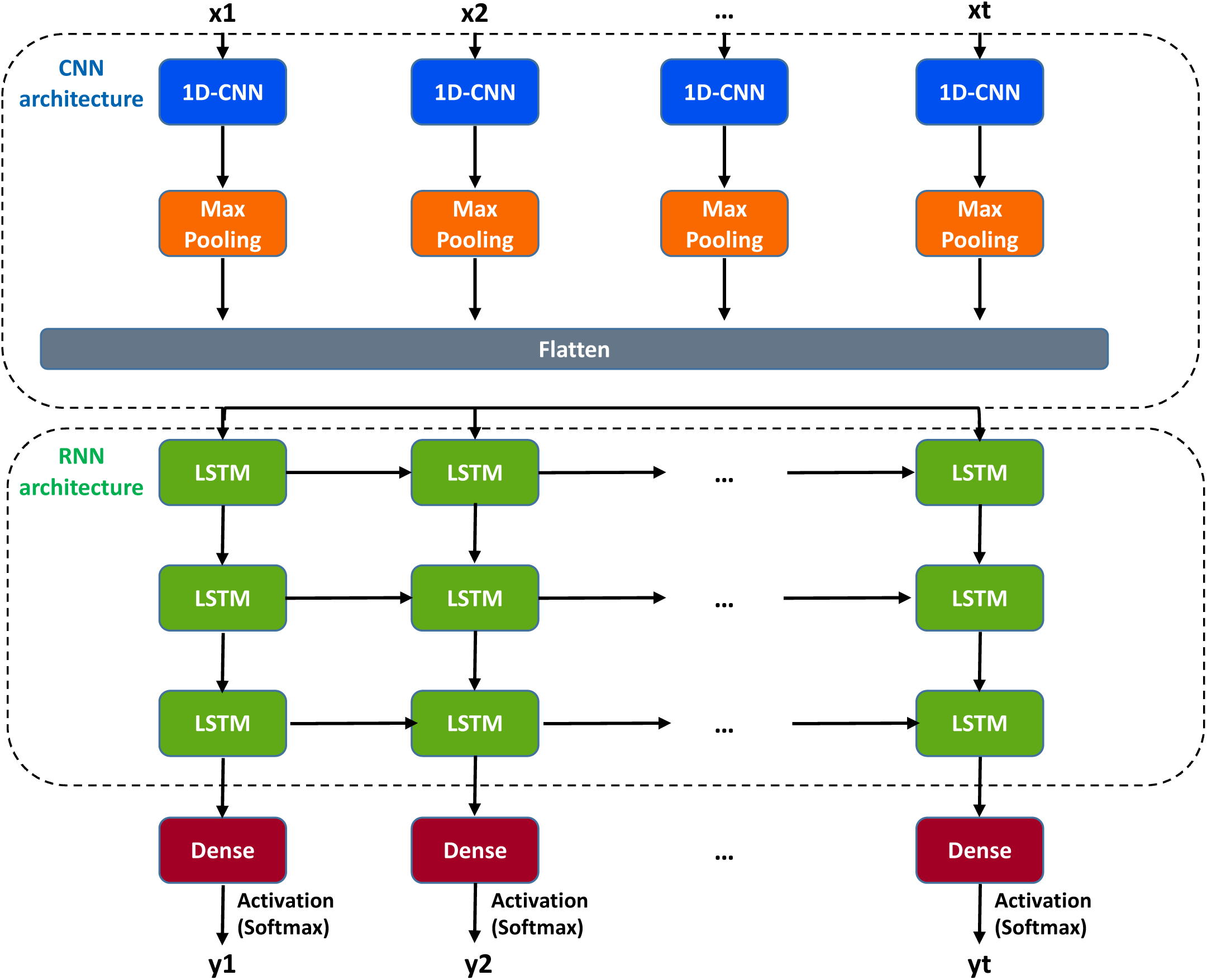
Deep-Channel model architecture. The input time series data was fed to the 1D Convolution layer (1D-CNN) which includes both 1D convolution layers, and max pooling layers. After this, data was flattened to the shape of the next network layer which is an LSTM. Three LSTM layers were stacked and each contains 256 LSTM units. Dropout layers were also appended to all LSTM layers with the value of 0.2 to reduce overfitting. This returned features from the stacked LSTM layers. The updated features are then forwarded to a regular dense layer with a SoftMax activation function giving an output representing the probability of each class (e.g. the probability × channels being open at each time step). In post-network processing, the most-likely number of channels open at a given time is calculated simply as the class with the highest probability at a given instant (Argmax).

**Fig.3.**
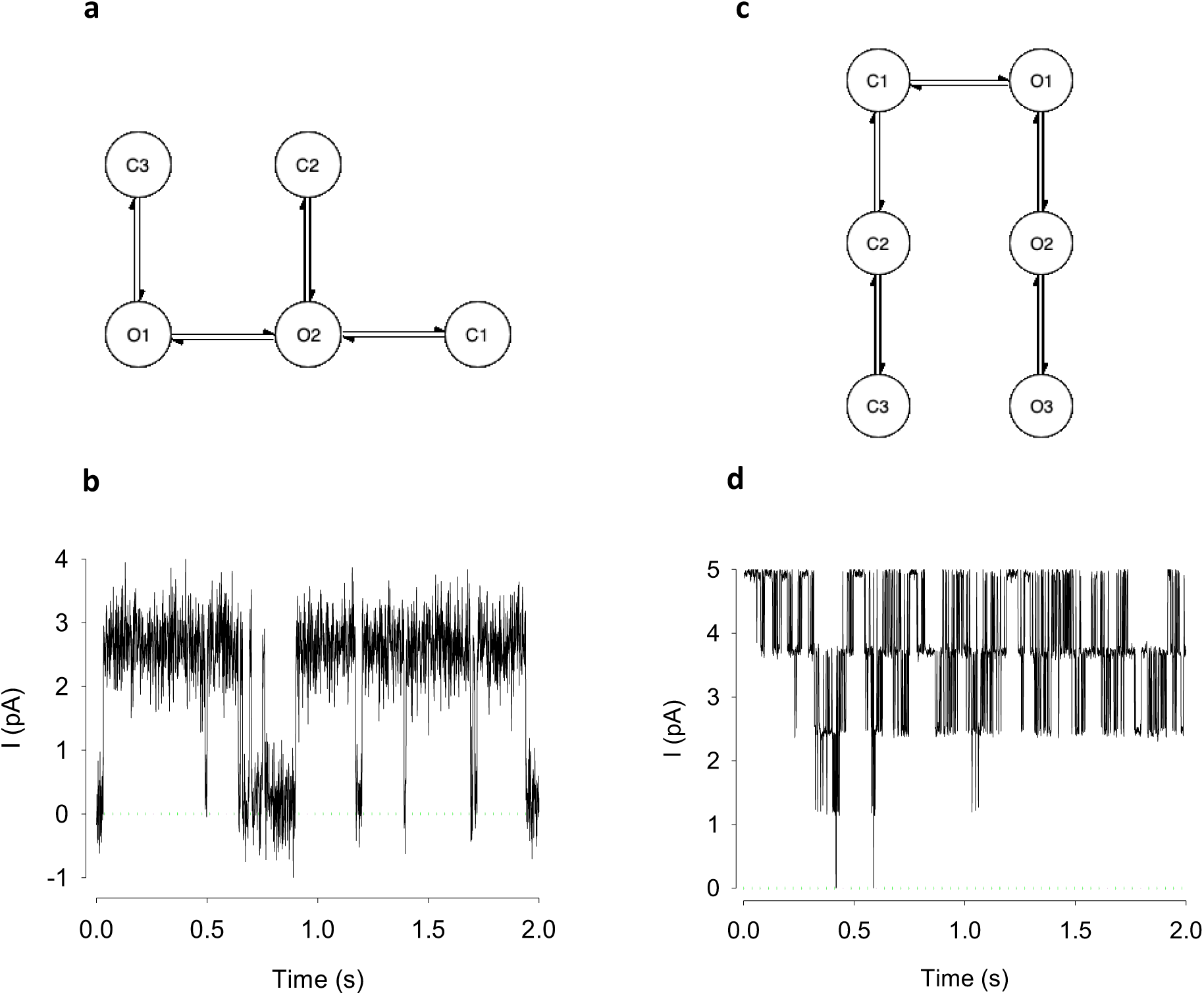
“Patch clamp” data was produced from two different stochastic models. **a,** and **c,** are the Markovian models used for simulation of ion channel data. Ion channels typically move between Markovian states that are either closed (zero conductance) or open (unitary conductance, *g*). The current passing when the channel is closed is zero (aside from recording offsets and artefacts), whereas when open the current (*i*) passing is given by *i* = *g* × *V*, where *V* is the driving potential (equilibrium potential for the conducting ion minus the membrane potential). In most cases there are several open and closed states (“O1”, “O2”, “O3”, or “C1”, “C2”, “C3” respectively). The central dogma of ion channel research is that the *g* will be the same for O1, O2 or O3. Although substates have been identified in some situations, these are beyond the scope of our current work. **a,** Model M1; the stochastic model from Davies et al^41^ and its output **b.** This model has a low open probability, and so the data is mostly a representation of zero or one channel open. **c,** Model M2; the stochastic model from^42^ and its output data **d**, since open probability is high the signal is largely composed of 3 or more channels simultaneously open.

**Fig. 4.**
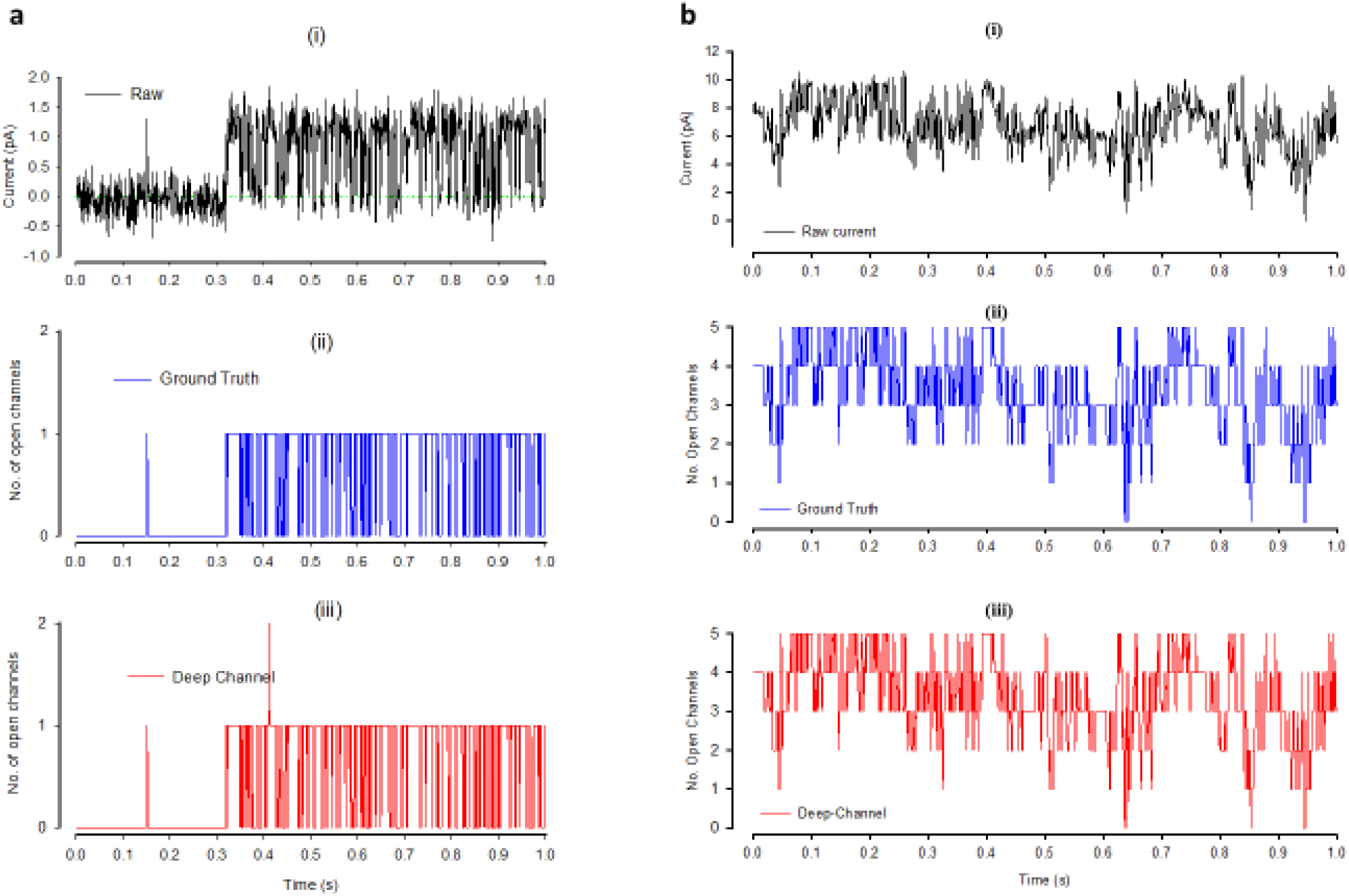
Qualitative performance Deep-Channel with previously unseen data. **a,** Representative example of Deep-Channel classification performance with low activity ion channels (data from model M1, Fig. 3a,b): **ai,** The raw semi-simulated ion channel event data (black). **aii,** The ground truth idealization/annotation labels (blue) from the raw data above in **ai**. **aiii,** The Deep-Channel predictions (red) for the raw data above **ai**. **b,** Representative example of Deep-Channel classification performance with 5 channels opening simultaneously (data from model M2, Fig. 3c,d). **bi,** The semi-simulated raw ion channel event data (black). **bii,** The ground truth idealization/annotation labels (blue) from the raw data above in **bi**. **biii,** The Deep-Channel label predictions (red) for the raw data above **bi**.

In datasets where channels had a low opening probability (i.e., from model M1), the data idealisation process becomes close to a binary detection problem (Fig. 4a), with ion channel events type closed or open (labels ‘0’ and ‘1’ respectively). In this classification, the ROC area under the curve (AUC) for both open and closed event detection exceeds 96% (Figure 5, Table 1). Full data for a representative example experiment is shown, with confusion matrix and ROC in Fig. 5a,b. Overall, in low open probability experiments, Deep-Channel returned a macro-F1 of 97.1±0.02% (standard deviation), *n*=10, whereas the SKM method in QuB resulted a macro-F1 of 95.5±0.025%, and 50% threshold method in QuB gave a macro-F1 of 84.7±0.05%, *n*=10.

**Fig. 5.**
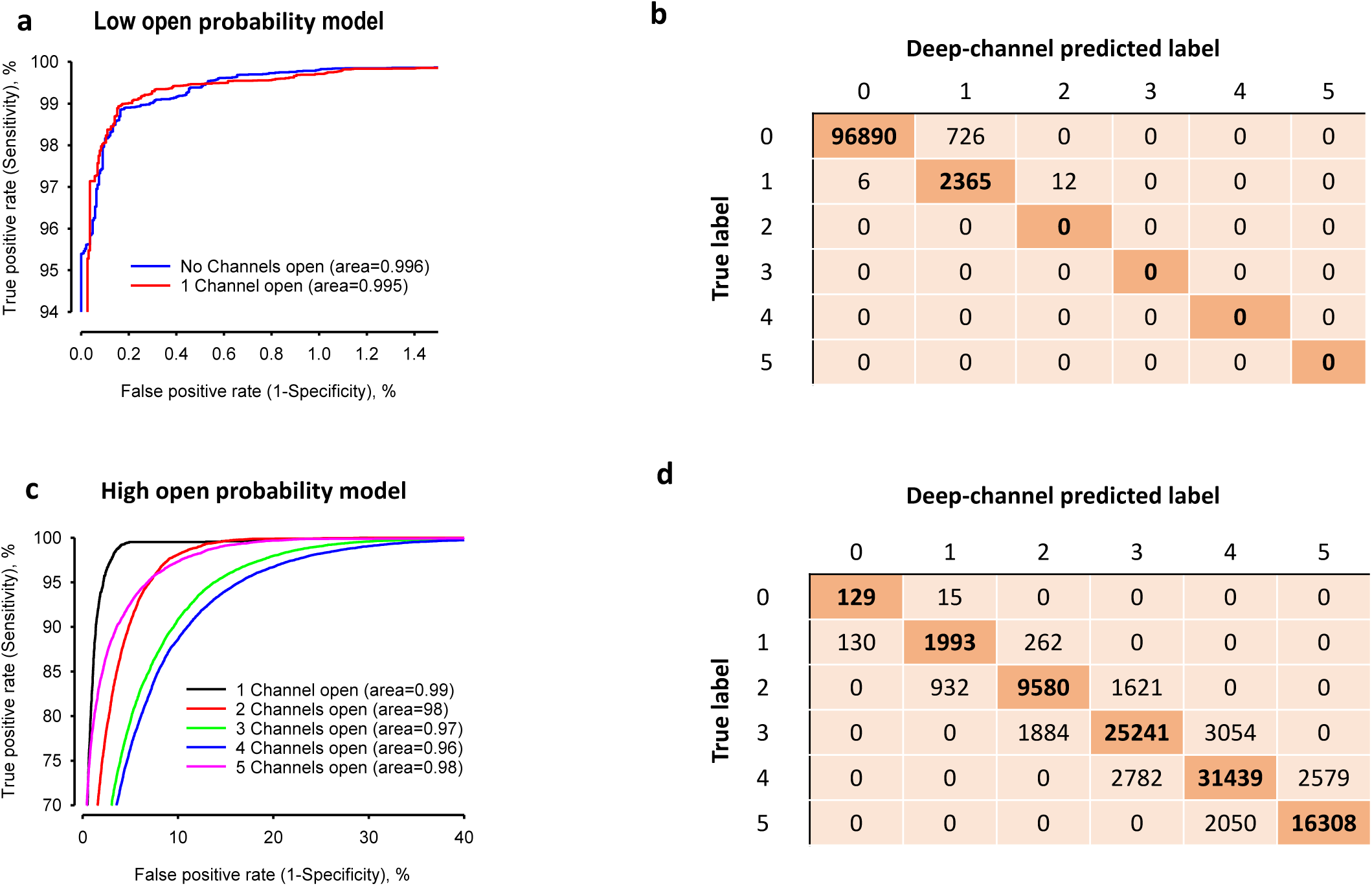
Quantitative performance of Deep-Channel with previously unseen data. **a,** Representative receiver operating characteristic (ROC) curve for ion channel event classification using the M1 stochastic gating model (Fig 3a.) and with only one channel present. **b,** The confusion matrix table for the example in **a**. Label 0 = no channels open, label 1= one channel open. Note that Deep-Channel is trained to recognise up to 5 channels opening at a time, however, with only one channel active at a time, the maximum ground truth class (True Label) is label 1 (one channel open). When analysing this simulated patch, Deep-Channel only made 12 (incorrect calls) of labels 2 to 5 (Deep-Channel predicted labels). **c,** Representative receiver operating characteristic (ROC) curve for ion channel event classification using the M2 stochastic gating model (Fig. 3) and with five channels present. Mean AUC are given in Table 1. **d,** The confusion matrix table for the example in **c**. Label 1= one channel open, label 2= 2 channels open etc. The mean ROC curve area under the curves (AUC) for all labels and all 17 experiments are given in Table 1.

**Table 1.** Deep-Channel ROC area under the curve (AUC) values achieved during 17 separate experiments, representative examples shown in Figure 5.

|  | Mean ROC AUC |  |
| --- | --- | --- |
| Number of Channels open simultaneously | low Popen data model M1, $n=10$<br>mean ± SD | High Popen data model M2, $n=7$<br>mean ± SD |
| 0 | 0.99±0.006 | 0.9997±0.0002 |
| 1 | 0.96±0.05 | 0.9973±0.0022 |
| 2 | - | 0.9917±0.0065 |
| 3 | - | 0.9824±0.0139 |
| 4 | - | 0.9787±0.0169 |
| 5 | - | 0.9937±0.0050 |

In cases where datasets included highly active channels (i.e., from model M2, Fig. 3c,d, Fig. 4b) this becomes a multi-class comparison problem and here, Deep-Channel outperformed both 50% threshold-crossing and SKM methods in QuB considerably. The Deep-channel macro-F1 for such events was 0.87±0.07 (standard deviation), *n*=7, however segmented-k means (SKM) macro-F1 in QuB, without manual baseline correction, dropped sharply to 0.57±0.15, and 50% threshold-crossing macro-F1 fell to 0.47±0.37 (Student’s paired t-test between methods, *p*=0.0052). An example ROC for high activity channel detection, and associated confusion matrix is shown in Fig. 5c,d. Frequently, in drug receptor or toxicity studies, biologists look for changes in open probability and so we also compared the open probability from manual threshold crossing, SKM and Deep-Channel against ground truth (fiducial). In the presence of several simultaneously opening channels in some quite noisy datasets with baseline drift, careful manual 50% threshold crossing and SKM sometimes essentially fail entirely, but Deep-Channel continues to be successful. For example, in some experiments with very noisy data threshold crossing open probability estimations were over 100% out and SKM detected only near half of the open events (~50% accuracy). Nevertheless, overall, there were highly significant correlations for both Deep-Channel vs ground truth (0.9998, 95% confidence intervals 0.9996 to 0.9999, *n=*17) and threshold crossing vs ground truth (0.95, 95% confidence intervals 0.87 to 0.98, *n=17*). In terms of speed, Deep-Channel consistently outperforms threshold crossing. Deep-channel analysed at the rate of approximately 10 seconds of data recording in under 4 seconds of computational time, whereas analysis time with threshold crossing in QuB was entirely dependent on the complexity of the signal.

Deep channel also proved robust to different levels of signal-to-noise ratios (SNR). For example, F1 scores in low, medium and high SNR levels are: low (SNR=5.35±2.18, F1=0.91±0.016), medium (SNR=12.74±3.65, F1=0.96±0.011), and high (SNR=60±4.52, F1=0.98±0.007).

### Biological ion channel data testing metrics

As stated earlier, a true Ground Truth is not possible with native ion channels signals recorded from biological membranes. However, with straightforward clear signals such as that shown in Figure 6, experts can idealise these data with supervised methods. We therefore chose a stretch of real data from^31^ including moderate level of noise and drift (Figure 6a). We then had 5 ion channel experts idealise these data. For each of the (approx.) 880,000 time-points we then took the mode of their binary idealisation value (0-closed or 1-open) to construct a “golden” dataset to use as an *effective* ground truth (Figure 6d & e). The consensus idealisation included 3241 openings. To check for inter-user agreement (Figure 6b) we calculated the over-all Fleiss Kappa^32^ implemented in R with the irr package, Fleiss Kappa was 0.953, with *p*≤1e-6. We then idealised this raw data (blinded from the “golden” dataset) with Deep-Channel and a range of other alternatives (Figure 6 f, g, h). The two alternatives we benchmark here are SKM using QuB^13^ and Minimum description length (MDL, using MatLab)^33^. Note that with Deep-Channel, there are no parameters to set and no post-processing. With SKM one needs to identify closed and open state levels and number of channels present. In the case of MDL there is no pre-processing necessary and no parameters to set, but the output is non-binary. Therefore, we ran a 50% event threshold crossing method on this to output final open and closed calls. These idealisations were all then compared to the “golden” dataset with a Cohen’s kappa agreement script, Table 2. Also, we fitted these data with a clustering and heatmap model in R, this allows one to visually compare the agreement at each timepoint.

**Fig. 6.**
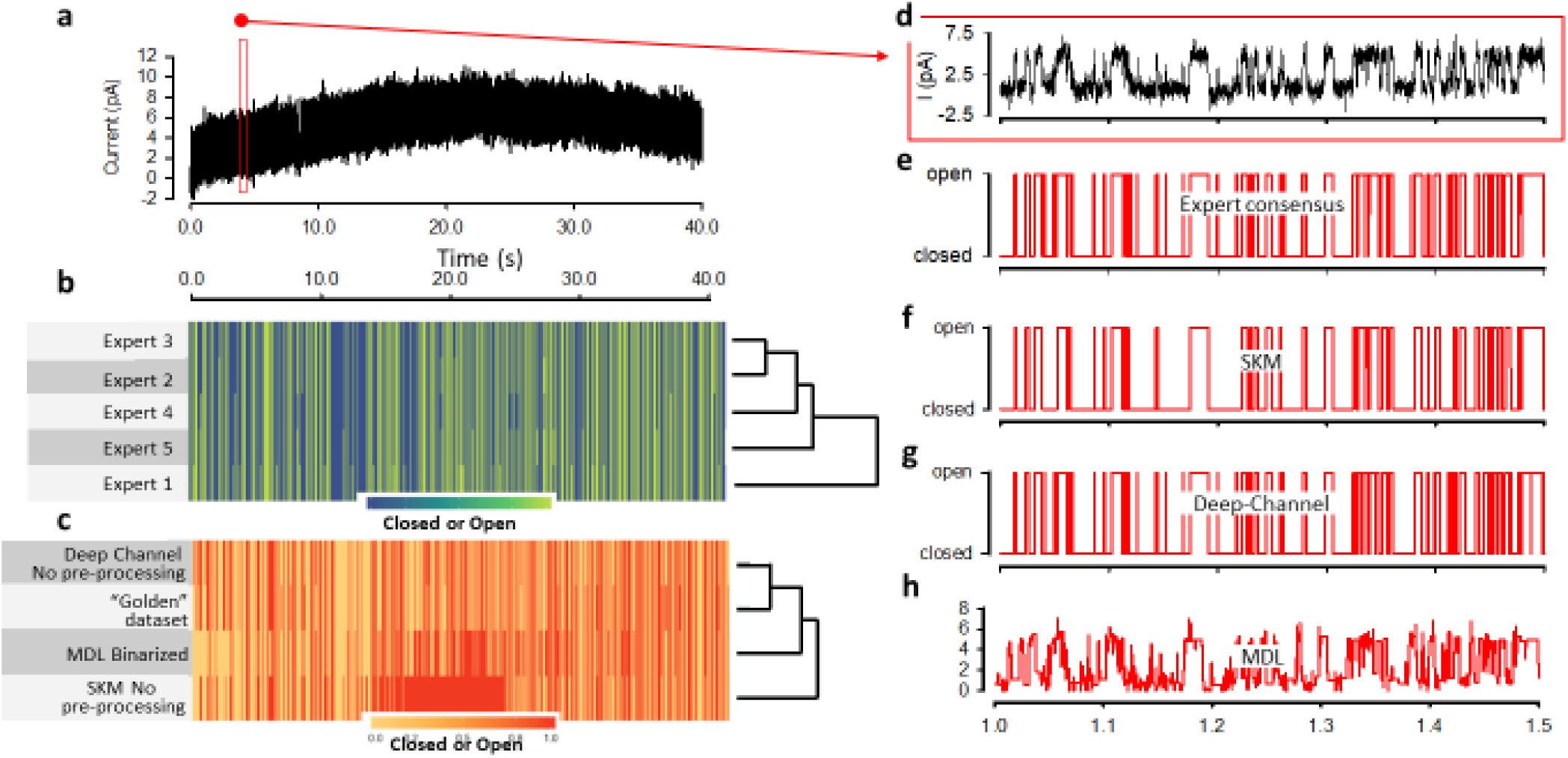
Performance of Deep-Channel with biological ion channel data. **a,** Single channel data from ^31^ with noise approximately 850,000 data points decimated by a factor of 50 for display only. **b,** The modal idealisation by 5 ion channel experts (“golden” idealisation). **c,** Agreement clustering and heatmap between “golden” dataset, MDL, Deep-Channel and SKM. **d,** A zoomed view of 500ms of raw biological data taken from (a). **e,** the expert consensus (modal) Idealisation. **f,** the idealisation output by SKM (after setting channel closed and open levels and setting the number of channels to 1), **g,** by Deep-Channel and (h) by MDL. Note that (c), (f), (g) and (h) were produced without baseline correction.

**Table 2.** The Cohen’s kappa agreement scores for automatic analyses, including Deep-channel, SKM, and MDL with golden dataset which is built by 5 different human experts using their existing software tools.

|  | Cohen's kappa score | 0.95% C.I. |
| --- | --- | --- |
| **MDL** | 0.766* | 0.7646 – 0.7674 |
| **SKM** | 0.6497** | 0.6481 - 0.6513 |
| **Deep-channel** | 0.9279 | 0.6481 - 0.6513 |

*\*MDL does not output an idealisation and so we binarized the MDL output using a 50% threshold crossing algorithm to allow comparison. \*\*SKM implemented in QuB has no true automatic function, one need to set the starting baseline, channel amplitude and number of channels before starting. We did not perform additional baseline correction or input baseline nodes for this comparison.*

## Discussion

Single molecule research, both FRET and patch-clamp electrophysiology provide high resolution data on the molecular state of proteins in real time, but their analyses are usually time consuming and require expert supervision. In this report, we demonstrate that a deep neural network, Deep-Channel, combining recurrent and convolutional layers can detect events in single channel patch-clamp data automatically. Deep-channel analysis is completely unsupervised and thus adds objectivity to single channel data analyses. With complex data, Deep-Channel also significantly outperforms traditional manual threshold crossing both in terms of *speed and accuracy*. We find this method works with very high accuracy across a variety of input datasets.

The most established single molecule method to observe single-channel gating is patch-clamp recording^2^. Its development led to the award of the Nobel Prize to Sakmann and Neher in 1991^34^ and the ability to observe single channels gate in real time validated the largely theoretical model of the action potential developed in the earlier Nobel Prize winning work of Hodgkin and Huxley^3^. Whilst the power and resolution of single-channel recording has never been questioned, it is well accepted to be a technically difficult technique to use practically since the data stream created requires laborious supervised analysis. In some cases, where several single channels gate simultaneously, it becomes impractical to analyse and data can be wasted. For practical purposes, drug screening etc, where subtle changes in channel activity could be crucial^5^, this means that the typical method is to measure bulk activity from a whole-cell simultaneously. Average current can be measured which is useful, but does not contain the detailed resolution that individual molecular recording has^14^. Furthermore, new technologies are emerging which can record ion channel data automatically^15, 35^, but whilst whole-cell currents are large enough to be analysed automatically, there are currently no solutions to do the same with single-channel events. Currently, it could be observed that automated patch-clamp apparatus are not used a great deal for single-channel studies and so automated analysis software are of little value, however, we feel that the reverse is true; this equipment is rarely used for single channel recording *because* no fully automated analysis exists. In this report we show that the latest machine learning methods, that of deep learning, including recurrent and convolutional neural network layers could address these limitations.

The fundamental limitation of applying deep learning to classification of biological data of all kinds is the prerequisite for training data. Deep learning is a form of supervised learning where during the training phase, the network must be taught at every single instant what the ground truth state is (it looks open, but is it really open or closed?). We considered two possible approaches to deal with this conundrum: (i) To collect data from easily analysable single molecule/patch clamp experiments and get a human expert to idealise this (classify or annotate it). This has two fundamental flaws that could be referred to as “Catch-22”. Firstly, if you train a network only to detect easy to analyse data, the output will be a network that can only detect easy-to-detect’’ events. Secondly, even then, if the events had to be human detected in the first place, it would mean that the final (trained) network would learn the same events as the human taught it; *and learn the same errors*. Analyses of ambiguous events would not tend toward detection with perfect accuracy, but inherit human biased errors. We therefore developed an alternative approach. (ii) Single channels gate in a stochastic, Markovian manner and therefore an unlimited number of idealised records can be simulated. This approach has been successfully applied before with other analyses development studies^12, 36^. The limitation is that there are inherent distortions and filtering that occur during collection of genuine data from a real analogue world. These can be imitated mathematically, but instead we used an entirely novel method of generating semi-synthetic training data; we played our idealised records out to a genuine patch-clamp amplifier using the dynamic-clamp approach^37^ and used an established analogue test cell (resistors and capacitors equivalent to a patch pipette and membrane). Our first data figure (Fig. 1b) shows the authenticity of this data and the approach. In summary, our novel methodology allows the creation of 100,000s training sets with noisy data in parallel to a ground truth idealisation. To conclude our work, we also compared deep channel performance against a simple “golden” idealisation by human supervised methods and two other existing methods. There are two obvious caveats with this approach, but we feel it is useful nonetheless. The first obvious caveat, is that in order for it to be possible to create the Golden datasets with human experts, it needed to be a very simple dataset with relatively few clear events. Secondly, it is not a ground truth. The small error between the Expert and Deep-Channel channels could be because the Experts were wrong rather than Deep-Channel. Bias, generally is discussed below, nevertheless the success of Deep-Channel in this experiment supports it’s potential for solving real ion channel idealisation problems.

Since our aim was to classify a time series, we developed a network with the combined power of both 1D-CNN layers and RNN (LSTM) units. Deep-Channel has a 1D-CNN at its core, but whilst ion channel activity is Markovian, the presence of both short and long duration underlying states means that it is important for a detection network to also be able to learn long-term dependencies across and so accuracy is improved with the LSTM (*see supplementary data*). Similar approaches combining RNNs and convolution layers have previously been applied to various analysis of biological gene sequences^38^ and cell detection in image classification^39^, but this is its first use for single-channel activity detection to our knowledge.

We used a number of metrics that are commonplace in machine learning and patch-clamp recording. Initially, to test the ability of Deep-Channel to detect events, we compared detected (predicted) idealised events against the fiducial idealised records. To compare against the human supervised methods, we analysed matching datasets with QuB and Deep-Channel and compared the summary parameter, open probability (*Po*) between them, together with F1 accuracies. Firstly, Deep-Channel was far quicker, to the degree that sometimes with complex multichannel data, manual analyses seems a stressful and near impossible task. With Deep-channel the complexity (within the datasets we used) made no difference to the speed or accuracy of performance. In terms of relative performance, note that whilst there was a strong correlation between open probabilities measured between Deep-Channel and threshold crossing, the F1-accuracy scores of the noisy data measured by threshold crossing fell of sharply. The significance of this is that whilst average open probability estimates from threshold crossing seem reasonable (some over estimates, some under-estimates cancelling out) the time-point by time-point accuracy *essential for kinetic analyses* is poor.

In the present work, we developed a method that works with channels of any size and kinetic distribution, but we did not include detection of multiple phenotypes or sub-conductances etc. A perceived problem, specifically with a machine learning model, is its generalisability. The concern is that the network would be good at detecting events in the exact dataset it was trained on, but fall short, when challenged with a quite different, but equally valid dataset; a problem known as over fitting. Furthermore, the heavy reliance on our model on training with synthetic data could lead to subtle and unexpected biases. In cell-attached mode, for example, the most feasible method for use in automated patch-clamp machines, there is often an asymmetry (relaxation) of larger events. This is shown to an extreme degree by Fenwick et al 1982^40^, but more subtle examples are often seen in native data. This could arrive from two situations, beyond the technical limitations of the headstage; for example, if the membrane potential changes during recording or if sufficient ion movement through the channel changes the ion driving force mid sojourn. Our training data did not see these types of anomalies. Furthermore, a subtle noise effect occurs with genuine ion channel events that experience patch clampers can see by eye; open channel noise as well as the well-known pink noise (1/f) of biological membranes. The noise level tends to increase during ion channel opening sojourns. To completely eliminate these or other biases long-term may be impossible, but potentially, mixing human annotated and synthetic data in an appropriate ratio may be one possible route. The goal would be to balance potential hidden biases from the synthetic data against the inevitable biases of human curation (humans will make human errors and be simply unable to label complex signals). Since perfect curation of simple datasets requires very simple datasets to annotate and is time consuming, potentially data augmentation could be used to build complex semi-synthetic datasets by building up layers of simpler data. Our tests against a Golden dataset produced by 5 ion channel experts found Deep-Channel to be remarkably good and it was a conservative test: (1) The SKM method required us to define the initial closed and open state levels, we did not do that in Deep-Channel. (2) The SKM method required us to define that there would be one channel present, we did not do that with Deep-Channel. It recognised that there was only 1 channel present, simply from the waveform. Currently, Deep-Channel can recognise up to 5 channels opening simultaneously.

We have demonstrated here the effectiveness of Deep-Channel, an artificial deep neural network to detect events in single molecule datasets, especially, but not exclusively patch-clamp data, but the potential for deep learning convolution/LSTM networks to tackle other problems cannot be overestimated.

## Methods

We develop a novel deep learning approach to automatically process large collections of single/multiple ion channel data series with detection of ion channel transition events, and re-construction of annotated idealised records. Datasets with pre-processing and analysis pipeline code will be made publicly available on GitHub (https://github.com/RichardBJ/Deep-Channel.git) including the model code to facilitate reproducibility. Fig. 1 shows an overall workflow and experimental design; creation of the digitised synthetic analogue datasets for developing a deep learning model, together with steps for training and testing (validating).

### Data description and dataset construction

Ion channel dwell-times were simulated using the method of Gillespie^43^ from published single channel models. Channels are assumed to follow a stochastic Markovian process and transition from one state to the next simulated by randomly sampling from a lifetime probability distribution calculated for each state. Authentic ‘electrophysiological’ noise was added to these events by passing the signal through a patch-clamp amplifier and recording it back to file with CED’s Signal software via an Axon electronic “model cell”. In some datasets additional drift was applied to the final data with Matlab. Two different stochastic gating models, (termed M1 and M2) were used to generate semi-synthetic ion channel data. M1 is a low open probability model from^41^ (Fig.3a, b), typically no more than one ion channel opens simultaneously. Model M2 is from^42, 44^ and has a much higher open probability (Fig. 3c, d), consequently up to 5 channels opened simultaneously and there are few instances of zero channels open. The source code for generating a combination of different single/multiple ion channel recordings is also given along with the publicly available datasets. Using this system, we can generate any number of training datasets with different parameters such as number of channels in the patch, number of open/close states, sampling frequency and temporal duration, based on published stochastic models. Fiducial, ground truth annotations for these datasets were produced simultaneously using MATLAB. Recordings were sampled at 10 kHz and each record had 10 seconds duration. To validate the Deep-Channel model, 6 different validation datasets were used: 3 datasets for single; and 3 datasets for multi-channel recordings. Datasets for training typically contained 10,000 subsets of 10 seconds each. Each dataset includes raw current data and ground truth state labels from the stochastic model, which we refer to as the idealisation. Within these training datasets, the third column is the fiducial record/ground truth and includes the class labels; ‘0’, ‘1’, ‘2’, ‘3’, ‘4’ and ‘5’. Each label indicates the instantaneous number(s) of open channels at a given time.

### Model background

LSTM (long short-term memory), is a type of RNN, the deep learning model architecture that is now widely adopted efficiently for time series forecasting with long-range dependencies. The major advantage of LSTM over RNN is its memory cell *c*_*t*_ which is computed by summing of the state information. This cell acts like a gate that activates or deactivates past information by several self-parametrized controlling gates including input, output and forget gates. As long as the input gate has a value of zero, then no information is allowed to access the cell. When a new input comes, its information is passed and summed to the cell if the input gate *i*_*t*_ is, in turn, activated. Ideally, the LSTM should learn to reset the memory cell information after it finishes processing a sequence and before starting a new sequence. This mechanism is dealt by forget gates *f*_*t*_ and the past cell content history *c*_*t*−1_ can be forgotten in this process and reset if the forget gate *f*_*t*_ is activated. Whether a cell output *c*_*t*_ will be passed to the final state *h*_*t*_ is further allowed by the output gate *o*_*t*_. The main innovation of using gating control mechanisms in LSTMs is that it ameliorates the vanishing gradient problem. This limitation of the general RNN model^27, 45^ is thus eliminated during forward and backward propagation periods. The key equations of an LSTM unit are shown in (1) below, where ‘∘’ denotes the Hadamard product:

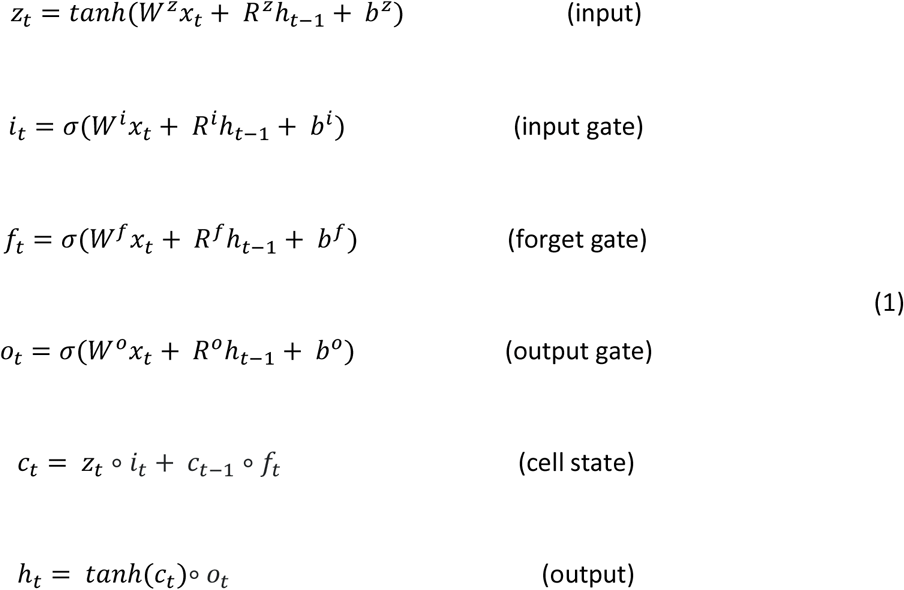

The *W*s* are input weights, the *R*s* are recurrent weights, *b*s* are the biases, *x*_*t*_ is denoted as the current input, and *h*_*t*−1_ is referred as the output from previous time step. The weighted inputs are accumulated and passed through tanh activation, resulting in *z*_*t*_. Multiple LSTMs can be stacked and temporarily combined with other types of deep learning architectures to form more complex structures. Such models have been applied to overcome previous sequence modelling problems^46^.

### Model development – Network architecture

Our Deep-Channel RCNN model was implemented in Keras with a Tensorflow backend^47^ using Python 3.6. Fig. 2 shows a graphical representation of the model architecture consisting of an input layer and 1D-convolution layer, a ReLU layer, a max pooling layer, a fully connected layer, 3-stacked LSTM with batch normalisation and drop out layers, and a final SoftMax^48^ output layer. For training; 3-dimensional data with ion channel current recordings (raw data), time steps (n=1), and features (n=1) served as input to the 1D convolution layer. This input layer feeds into a temporal convolution layer to investigate frame-level features. Afterwards, the output of the flattened convolutional layer is fed to the 3-stacked LSTM layers. Finally, the network feeds into one dense neuron with a SoftMaxactivation function, outputting the probability of a given channel level. This combined 1D convolution LSTM (RCNN) model was then saved as a hierarchical data format file HDF5 to allow automatic detection in other datasets without need to retrain. HDF5 files, including trained tensor weighting, are also available via our GitHub site.

### Class Imbalance

Typical real-world ion channel time series data are usually inherently class imbalanced (see for example, the confusion matrices in Fig. 5); if only one channel is present, there could be very few openings or very few closures. If there is more than one channel in the patch, the number of channels open at any one instant will be distributed binomially. This increases the volume of data required to train the network. To address this, where we looked at highly active channels, our training process used an oversampling of the minority classes, rearranging datasets evenly using the synthetic minority oversampling technique^49^, implemented in the Python Imbalanced-Learn library^50^. SMOTE adds the over-samples to the end of the end of the data record, but for training purposes we shuffled those back into the body of the data.

### 1D-Convolutional Layer

The 1D Convolution layer (1D-CNN) step consists of 1D-CNN, rectified linear unit (ReLU) layer^51^ and max pooling layer. We used 64 filters and ReLU was applied as an activation function. After that, the max-pooling layer was added to each output to extract a representative value. Finally, data was flattened to allow input to the next network layer, an LSTM.

### RNN-LSTM

Three LSTM layers were stacked and each contains 256 LSTM units with ReLU activation functions. In the next step, a batch normalization (BN)^52^ was applied to standardize the inputs, meaning the mean will be close to 0, and the standard deviation close to 1, hence the training of the model is accelerated. Dropout layers were also appended to all RNN-LSTM layers with the value of 0.2 to reduce overfitting^53^. The returned features from the stacked RNN-LSTM layers were then fed into a flatten layer to have a suitable shape for the final layer. The updated features are then forwarded to a Dense output layer with a SoftMax activation function, in which output features one dense neuron PER potential channel level (zero to maximum channels). In our current model this maximum value is 5 channels. The output is the probability of each given class (e.g. the probability of × channels being open at each timepoint). In order to calculate the final classification, we take the class with maximum value probability at each instant.

### Model training

In the model training stage, once the probability values are calculated, errors between the predicted values and true values were calculated with a sparse categorical cross entropy as a loss function. To optimize the loss value, stochastic gradient descent was applied as an optimizer with an initial learning rate of 0.001, momentum of 0.9, and the size of a mini-batch was set to between 256 and 2048 depending on the model. A learning rate decaying strategy was employed to the model to yield better performance. Based on this strategy, the learning rate (initially is 0.001) was decayed at each 10^th^ epoch with decaying factor 0.01 of learning rate. The proposed Deep-Channel model was trained for 50 epochs. In the case of the training data an 80% -train and 20%-test split was performed.

### Performance Metrics

One of the clearest quality indicators of a classification deep learning method is the confusion matrix, sometimes called as contingency table^54^. This consists of four distinctive parameters, which are true positives *(TP)*, false positives *(FP)*, false negatives *(FN)*, and true negatives *(TN).* If the model’s output accurately predicts the ion channel event, it is considered as TP. On the other hand, it is indicated as FP if the model incorrectly detected an ion channel event when there is no a channel opening. If the model output misses an ion channel event activity, then it is computed as FN. These metrics are used in calculation of evaluation metrics such as precision (positive predicted value), recall (sensitivity), and F-score as described below in (3), (4), and (5) respectively:

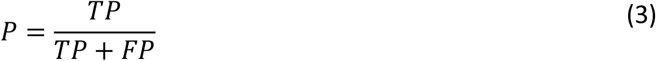

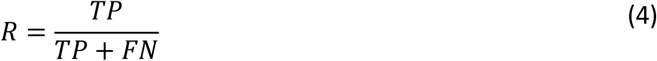

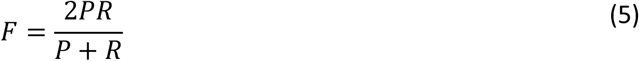

where *P*, *R*, and *F* denote precision, recall and F-score, respectively. In addition, area under curve (AUC) and receiver operating characteristic (ROC) parameters are efficiently used to visualize the model performance in classification problems. The ROC shows the probability relations between true positive rate (sensitivity-recall), and false positive rate (1-specificity), while AUC represents a measure of the separability between classes.

As an additional metric, more familiar to electrophysiologists we also calculated the open probability (Po), and compared this metric between Deep-Channel, a traditional software package (QuB) and the ground truth. The equation for open probability is given in (equation 6) as follows:

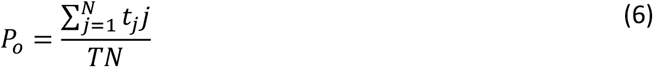

where *T* denotes total time, *N* is defined as numbers of channels in the patch, and *tj* is referred to the time spent with *j* channels open^55^. Since true numbers of channels in a patch is always an unknown parameter this was estimated as the maximum number of simultaneous openings.

### Computer hardware

In this work, we trained the Deep-Channel model on a workstation with an Nvidia Geforce GTX 1080Ti and 32GB RAM. The entire process of the proposed model including training, validating, evaluating, and visualizing process was employed within Python Spyder 3.6. Speed estimations were made with a similar PC, but without a GPU.

### Statistics

Statistical analyses were performed with R-studio, means are quoted with standard deviations, *n* is the number of experiments.

## Data availability

Datasets with pre-processing and analysis pipeline code will be made publicly available on GitHub (https://github.com/RichardBJ/Deep-Channel.git)

## Code availability

All code will be available on GitHub (https://github.com/RichardBJ/Deep-Channel.git)

## Acknowledgements

This work was funded by a BBSRC Transformative Resources Development Fund award.

## Author Contributions

RBJ, YZ and FC designed the study.

RBJ and NC wrote the code, NC prepared all the datasets.

RBJ, FMOB, CD, SB, NC, RDR contributed to the experimental work and the analysis and interpretation.

RBJ, NC, YZ, FC, FMOB drafted the initial manuscript.

All authors critically evaluated, edited and approved the final manuscript.

## Competing Interests

All authors declare that they have no competing interests.

## Notes

#### Summary of Updates

Up dates suggested by reviewers, but still not accepted!

https://github.com/RichardBJ/Deep-Channel

## References

1. Hamill, O.P., Marty, A., Neher, E., Sakmann, B. & Sigworth, F.J. Improved Patch-Clamp Techniques for High-Resolution Current Recording from Cells and Cell-Free Membrane Patches. Pflugers Archiv-European Journal of Physiology 391, 85–100 (1981).

2. Neher, E. & Sakmann, B. Single-channel currents recorded from membrane of denervated frog muscle fibres. Nature 260, 799–802 (1976).

3. Hodgkin, A.L. & Huxley, A.F. A Quantitative Description of Membrane Current and Its Application to Conduction and Excitation in Nerve. Journal of Physiology-London 117, 500–544 (1952).

4. Abdul Kadir, L., Stacey, M. & Barrett-Jolley, R. Emerging Roles of the Membrane Potential: Action Beyond the Action Potential. Front Physiol 9, 1661 (2018).

5. Lehmann-Horn, F. & Jurkat-Rott, K. Voltage-gated ion channels and hereditary disease. Physiological Reviews 79, 1317–1372 (1999).

6. Colquhoun, D. & Sigworth, F. in Single-channel recording 483–587 (Springer, 1995).

7. Qin, F., Auerbach, A. & Sachs, F. A direct optimization approach to hidden Markov modeling for single channel kinetics. Biophysical Journal 79, 1915–1927 (2000).

8. O’Brien, F. et al. Enhanced activity of multiple TRIC-B channels: an endoplasmic reticulum/sarcoplasmic reticulum mechanism to boost counterion currents. J Physiol 597, 2691–2705 (2019).

9. Ha, T. Single-molecule fluorescence resonance energy transfer. Methods 25, 78–86 (2001).

10. Blanco, M. & Walter, N.G. Analysis of complex single-molecule FRET time trajectories. Methods Enzymol 472, 153–178 (2010).

11. Juette, M.F. et al. Single-molecule imaging of non-equilibrium molecular ensembles on the millisecond timescale. Nat Methods 13, 341–344 (2016).

12. Colquhoun, D., Hatton, C.J. & Hawkes, A.G. The quality of maximum likelihood estimates of ion channel rate constants. Journal of Physiology-London 547, 699–728 (2003).

13. Nicolai, C. & Sachs, F. Solving ion channel kinetics with the QuB software. Biophysical Reviews and Letters 8, 191–211 (2013).

14. Sivilotti, L. & Colquhoun, D. In praise of single channel kinetics. J Gen Physiol 148, 79–88 (2016).

15. Dunlop, J., Bowlby, M., Peri, R., Vasilyev, D. & Arias, R. High-throughput electrophysiology: an emerging paradigm for ion-channel screening and physiology. Nat Rev Drug Discov 7, 358–368 (2008).

16. LeCun, Y., Bengio, Y. & Hinton, G. Deep learning. Nature 521, 436–444 (2015).

17. Boza, V., Brejova, B. & Vinar, T. DeepNano: Deep recurrent neural networks for base calling in MinION nanopore reads. PLoS One 12, e0178751 (2017).

18. Albrecht, T., Slabaugh, G., Alonso, E. & Al-Arif, S. Deep learning for single-molecule science. Nanotechnology 28, 423001 (2017).

19. Angermueller, C., Parnamaa, T., Parts, L. & Stegle, O. Deep learning for computational biology. Mol Syst Biol 12, 878 (2016).

20. Krizhevsky, A., Sutskever, I. & Hinton, G.E. in Advances in neural information processing systems 1097–1105 (2012).

21. Pratt, H. et al. Automatic Detection and Distinction of Retinal Vessel Bifurcations and Crossings in Colour Fundus Photography. Journal of Imaging 4 (2018).

22. Al-Bander, B., Al-Nuaimy, W., Williams, B.M. & Zheng, Y.L. Multiscale sequential convolutional neural networks for simultaneous detection of fovea and optic disc. Biomedical Signal Processing and Control 40, 91–101 (2018).

23. Misiunas, K., Ermann, N. & Keyser, U.F. QuipuNet: Convolutional Neural Network for Single-Molecule Nanopore Sensing. Nano Lett 18, 4040–4045 (2018).

24. Azizi, S. et al. Deep Recurrent Neural Networks for Prostate Cancer Detection: Analysis of Temporal Enhanced Ultrasound. IEEE Trans Med Imaging 37, 2695–2703 (2018).

25. Tang, D., Qin, B. & Liu, T. in Proceedings of the 2015 conference on empirical methods in natural language processing 1422–1432 (2015).

26. Hochreiter, S. The vanishing gradient problem during learning recurrent neural nets and problem solutions. Int J Uncertain Fuzz 6, 107–116 (1998).

27. Hochreiter, S. & Schmidhuber, J. Long short-term memory. Neural Computation 9, 1735–1780 (1997).

28. Graves, A., Mohamed, A.-r. & Hinton, G. in 2013 IEEE international conference on acoustics, speech and signal processing 6645–6649 (IEEE, 2013).

29. Sutskever, I., Vinyals, O. & Le, Q.V. Sequence to Sequence Learning with Neural Networks. Adv Neur In 27 (2014).

30. Lipton, Z.C., Berkowitz, J. & Elkan, C. A critical review of recurrent neural networks for sequence learning. arXiv preprint arXiv:1506.00019 (2015).

31. Mobasheri, A. et al. Characterization of a stretch-activated potassium channel in chondrocytes. Journal of Cellular Physiology 223, 511–518 (2010).

32. Fleiss, J.L. Measuring Nominal Scale Agreement among Many Raters. Psychol Bull 76, 378–& (1971).

33. Gnanasambandam, R. et al. Unsupervised Idealization of Ion Channel Recordings by Minimum Description Length: Application to Human PIEZO1-Channels. Front Neuroinform 11, 31 (2017).

34. Aldhous, P. Nobel prize. Patch clamp brings honour. Nature 353, 487 (1991).

35. Yajuan, X., Xin, L. & Zhiyuan, L. A comparison of the performance and application differences between manual and automated patch-clamp techniques. Curr Chem Genomics 6, 87–92 (2012).

36. Mukhtasimova, N., daCosta, C.J. & Sine, S.M. Improved resolution of single channel dwell times reveals mechanisms of binding, priming, and gating in muscle AChR. J Gen Physiol 148, 43–63 (2016).

37. Sharp, A.A., O’Neil, M.B., Abbott, L.F. & Marder, E. Dynamic clamp: computer-generated conductances in real neurons. J Neurophysiol 69, 992–995 (1993).

38. Lanchantin, J., Singh, R., Wang, B.L. & Qi, Y.J. Deep Motif Dashboard: Visualizing and Understanding Genomic Sequences Using Deep Neural Networks. Biocomput-Pac Sym, 254–265 (2017).

39. Kitrungrotsakul, T. et al. in ICASSP 2019-2019 IEEE International Conference on Acoustics, Speech and Signal Processing (ICASSP) 1239–1243 (IEEE, 2019).

40. Fenwick, E.M., Marty, A. & Neher, E. A patch-clamp study of bovine chromaffin cells and of their sensitivity to acetylcholine. J Physiol 331, 577–597 (1982).

41. Davies, L.M., Purves, G.I., Barrett-Jolley, R. & Dart, C. Interaction with caveolin-1 modulates vascular ATP-sensitive potassium (K(ATP)) channel activity. Journal of Physiology-London 588, 3254–3265 (2010).

42. O’Brien, F. & Barrett-Jolley, R. CVS role of TRPV: from single channels to HRV assessment with Artificial Intelligence. FASEB J 32, 732.736 (2018).

43. Gillespie, D.T. Exact Stochastic Simulation of Coupled Chemical-Reactions. J Phys Chem-Us 81, 2340–2361 (1977).

44. Feetham, C.H., Nunn, N., Lewis, R., Dart, C. & Barrett-Jolley, R. TRPV4 and KCa ion channels functionally couple as osmosensors in the paraventricular nucleus. Br J Pharmacol 172, 1753–1768 (2015).

45. Karpathy, A. & Li, F.F. Deep Visual-Semantic Alignments for Generating Image Descriptions. Proc Cvpr Ieee, 3128–3137 (2015).

46. Shi, X.J. et al. Convolutional LSTM Network: A Machine Learning Approach for Precipitation Nowcasting. Advances in Neural Information Processing Systems 28 (Nips 2015) 28 (2015).

47. Chollet, F. Deep Learning with Python. (Manning Publications Co., 2017).

48. Goodfellow, I., Bengio, Y. & Courville, A. Deep Learning. Adapt Comput Mach Le, 1–775 (2016).

49. Chawla, N.V., Bowyer, K.W., Hall, L.O. & Kegelmeyer, W.P. SMOTE: Synthetic minority over-sampling technique. J Artif Intell Res 16, 321–357 (2002).

50. Lemaitre, G., Nogueira, F. & Aridas, C.K. Imbalanced-learn: A Python Toolbox to Tackle the Curse of Imbalanced Datasets in Machine Learning. J Mach Learn Res 18 (2017).

51. Nair, V. & Hinton, G.E. in Proceedings of the 27th international conference on machine learning (ICML-10) 807–814 (2010).

52. Ioffe, S. & Szegedy, C. Batch normalization: Accelerating deep network training by reducing internal covariate shift. arXiv preprint arXiv:1502.03167 (2015).

53. Srivastava, N., Hinton, G., Krizhevsky, A., Sutskever, I. & Salakhutdinov, R. Dropout: A Simple Way to Prevent Neural Networks from Overfitting. J Mach Learn Res 15, 1929–1958 (2014).

54. Story, M. & Congalton, R.G. Accuracy assessment: a user’s perspective. Photogrammetric Engineering and remote sensing 52, 397–399 (1986).

55. Barrett-Jolley, R., Comtois, A., Davies, N.W., Stanfield, P.R. & Standen, N.B. Effect of adenosine and intracellular GTP on K-ATP channels of mammalian skeletal muscle. Journal of Membrane Biology 152, 111–116 (1996).

